# Group Behaviors Foster Zebrafish Foraging Through Social Interaction

**DOI:** 10.64898/2026.07.16.738874

**Authors:** Peishi Wang, Mingyang Chen, Bo Li

## Abstract

Group living enhances foraging efficiency, yet whether this benefit extends across developmental stages and how social coordination versus emotional buffering contribute remain unresolved. Using zebrafish (larvae 10 dpf, adults 80 dpf) in a Y-maze foraging task with high-resolution tracking and mirror experiments, we show that grouping reduces foraging latency in adults, but not larvae. Adults form coordinated groups (high polarization, small inter-individual distances), whereas larvae form loose aggregations. Adults, but not larvae, exhibit reduced freezing (social buffering) when grouped. Mirror-presented visual cues suffice to enhance foraging via emotional calming—reducing freezing and stabilizing speed—yet disrupt real-group coordination. Our results reveal a dual mechanism— instrumental coordination and emotional buffering—by which group behavior promotes foraging. Both mechanisms are developmentally gated, emerging only after maturation, offering insights into the ontogeny of social competence and practical implications for aquaculture and conservation.

## Introduction

Animals frequently aggregate in highly organized collectives—bird flocks, fish schools, insect swarms, and ungulate herds—that emerge from local interactions without central control [1,2]. These collective patterns carry profound ecological significance, prominently enhancing survival through reduced predation risk, improved foraging efficiency, and rapid information transfer [3–9]. Foraging, a fundamental activity bridging physiology and environment, is well described by optimal foraging theory and the marginal value theorem, which predict strategies that maximize net energy gain and govern patch-leaving decisions [10–13]. Foraging success is tightly linked to social interactions, where individuals rely on visual attention and neural processing to detect and integrate social cues, facilitating shared resource discovery and social learning about food and threats [14]. Yet group living also imposes costs, including resource competition, disease transmission, and social conflict, and understanding how this benefit–cost balance is shaped—especially across developmental stages—remains a central challenge in behavioral ecology [15,16].

While group foraging advantages are well documented, it remains unclear whether benefits stem from social coordination (aligned movement and cohesion) or social buffering (stress reduction via conspecific presence), whether these mechanisms are accessible across development, and whether visual social cues alone suffice. Foraging is suppressed by stress, evident as freezing [17,18]; social buffering, linked to oxytocinergic and amygdala–prefrontal circuitry, mitigates stress in mammals during specific developmental windows [19–22], yet its developmental constraints in zebrafish are unexplored [23]. Mirror assays separate the emotional component of visual companionship from interactive coordination [24–26], but few studies have applied this to foraging or examined age dependence. Key factors affecting foraging thus include stress, social interaction, visual attention, and brain circuit maturation. Cross-species comparisons reveal a conserved critical window for social competence requiring prefrontal–amygdala maturation [27–29], and dynamic oxytocin receptor expression in zebrafish implies larvae may lack the neural circuits to extract foraging benefits from group living [30,31]. A systematic understanding of how social mechanisms facilitating foraging emerge across development therefore represents a critical gap.

Zebrafish (Danio rerio) are a powerful model for studying social behavior and development due to their sociality, shoaling/schooling, multi-sensory communication (vision and lateral line), rapid generation time, optical transparency of larvae for neural observation, and rich genetic tools for targeted manipulations [32–37]. Modern tracking and machine learning enable extraction of precise kinematic parameters—speed, polarization, inter-individual distance, and freezing—to characterize group dynamics at high resolution [38–41]. However, most studies focus on adults, leaving a gap in how developmental stage shapes group-living outcomes [42,43]. Our recent work identifies a key developmental window for the establishment of collective behavior [44], raising an intriguing question whether group behavior affect foraging. It is documented that adult group foraging benefits [42], whether advantages derive from social coordination or social buffering, and how these mechanisms emerge developmentally, remain unknown.

To address this, we systematically compared foraging performance, kinematics, and stress behaviors between larvae (10 dpf) and adult (80 dpf) zebrafish under solitary, group, and mirror conditions in a Y-maze foraging task. We asked three central questions: does group foraging advantage persist across development; if age-dependent, does it stem from social coordination, social buffering, or both; and are visual social cues alone sufficient, and if so, by what mechanism? We found that group advantages are strictly developmentally gated: adults benefited strongly, whereas larvae showed no improvement over solitary fish. This adult advantage reflects a dual mechanism—enhanced social coordination (higher polarization, lower inter-individual distance) and robust social buffering (reduced freezing)—both absent in larvae. Mirror cues alone partially improved foraging through emotional calming, but could not reproduce the coordinated interactions of a real group. These findings elucidate the ontogeny of social competence in zebrafish and provide a mechanistic framework for how group living promotes individual fitness.

### Group housing enhanced foraging efficiency in adults, but conferred no such benefit on larvae

To assess the effect of group behavior on foraging efficiency in zebrafish, we conducted behavioral tests using a Y-maze apparatus (**Fig. 1A**, **Movie 1-3**). We compared the foraging latency—defined as the time interval from entering the experimental arena to first feeding—between solitary (n = 1) and group (n = 10) conditions in both larvae and adults (**Fig. 1B**). Consistent with our initial hypothesis, statistical analysis of the experimental data (**Fig. 1C**) revealed that, in adults, the foraging latency was significantly shorter in the group condition compared to the solitary condition (n = 100, p = 0.0011), indicating that group behavior indeed enhances foraging efficiency.

**Figure 1.**
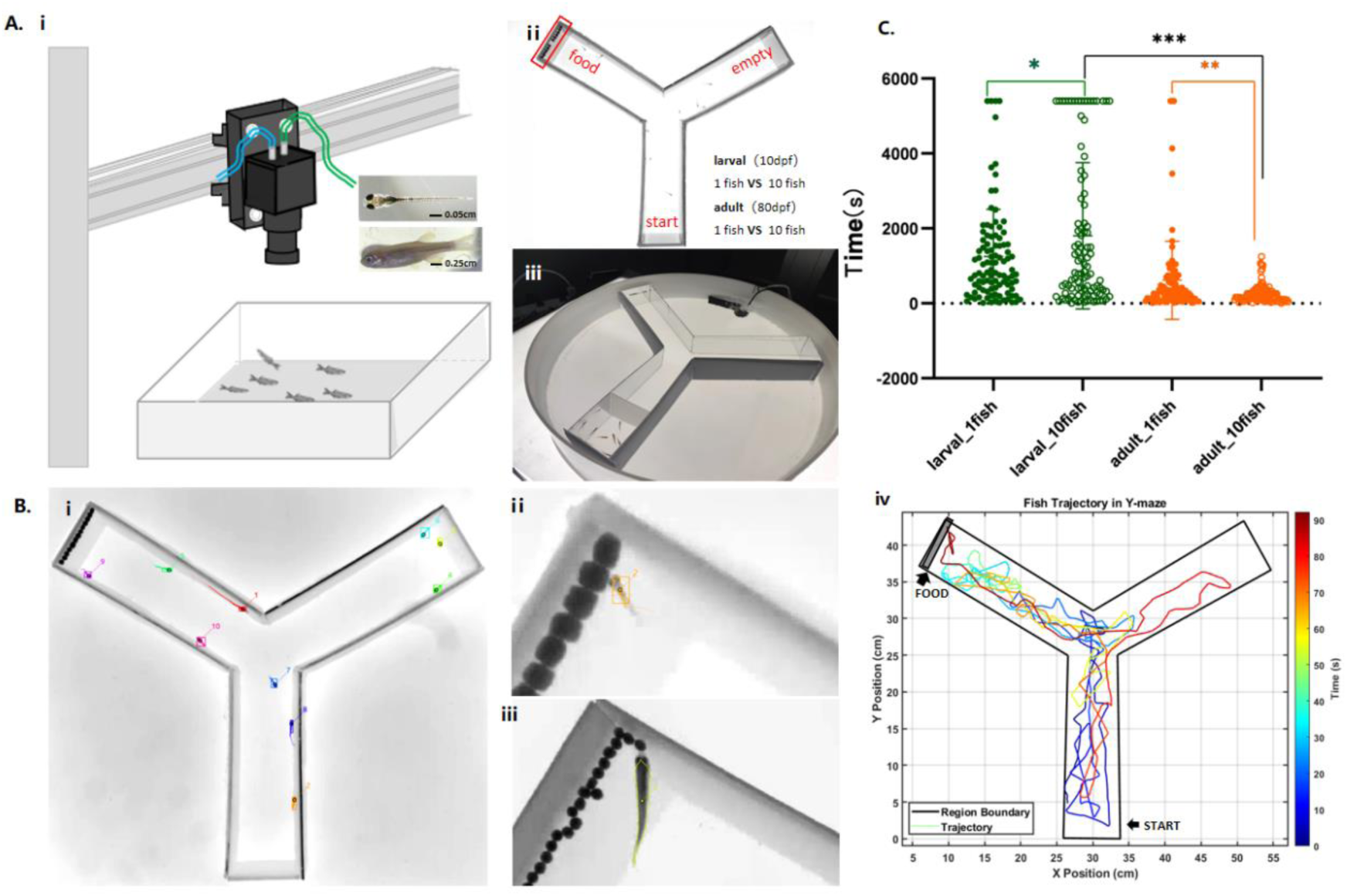
Overview. **(A) Experimental apparatus. Panel (ⅰ)** shows a schematic diagram of the experimental setup. An industrial camera was mounted perpendicularly above the water surface, and a backlight panel placed at the bottom of the tank provided illumination. The effective experimental arena consisted of a Y-maze, with a top-view schematic shown in panel **(ⅱ).** During experiments, food was placed at the end of one arm, and the other arm served as the starting arm. Two tank sizes were designed based on the developmental stage of the experimental subjects. For larval experiments (10 dpf, body length approximately 0.5 cm), the Y-maze arm length was 5 cm, with a channel width of 1.6 cm and a height of 1.6 cm. For adult experiments (80 dpf, body length approximately 2.5 cm), the Y-maze arm length was 25 cm, with a channel width of 8 cm and a height of 8 cm. Panel **(ⅲ)** shows a photograph of the adult experimental tank. **(B) Data extraction. (ⅰ)** Trajectory tracking was performed using idtrackerai. Panels **(ⅱ)** and **(ⅲ)** show photographs of larval and adult foraging moments, respectively. Panel **(ⅳ)** displays representative movement trajectories of fish within the experimental arena, with colors representing different time points as indicated by the jet colormap on the right: colors ranging from blue (start of trial) to red (end of trial). **(C) Foraging time analysis.** T-test (two-tailed one-sample t-test) was used to examine correlations between different movement parameters across experimental groups. Solid green dots represent the solitary larvae group, hollow green dots represent the larvae group, and solid orange dots represent the solitary adult group. larvae solitary vs. larvae group: p = 0.0185, p < 0.05, *; adult solitary vs. adult group: p = 0.0011, p < 0.05, **; larvae solitary vs. adult solitary: p = 0.0001, p < 0.05, ***; larvae group vs. adult group: p < 0.0001, ****.

Notably, however, this group advantage completely disappeared in larvae. Interestingly, foraging latency in the group condition was even slightly longer than that in the solitary condition, although the difference was not statistically significant (n = 100, p = 0.0185). These results demonstrate that the facilitative effect of grouping on foraging efficiency is strongly developmentally dependent, suggesting that the maturation of social coordination may require a specific developmental time window. This raises the question: what mechanisms enable adults, but not larvae, to gain a foraging advantage from social context? We sought to address this question by examining both the kinetic and behavioral aspects of group dynamics.

### The kinematic basis of group foraging advantage—Group coordination of adult fish

To uncover the kinematic mechanisms underlying the age-dependent group foraging advantage, we first analyzed movement trajectories across experimental conditions. **Fig. 2A** shows representative paths of a solitary larvae (ⅰ), a larvae group (ⅱ), a solitary adult (ⅲ), and an adult group (ⅳ), with color coding from blue to red indicating progression from the start to the end of the trial. Qualitative inspection of the trajectories revealed no overt differences across conditions: fish in all four groups exhibited extensive exploration of the experimental arena, repeatedly entering the food arm and approaching the food patch multiple times before finally feeding (**Fig. 2A**). This suggests that, regardless of age or social context, zebrafish engage in exploratory behavior upon entering a novel environment. Notably, some individual variation in this exploratory pattern was observed—a few fish swam directly toward the food upon detection and fed immediately (**Fig. S1**)—but overall, the spatial utilization patterns were similar across groups.

**Figure 2.**
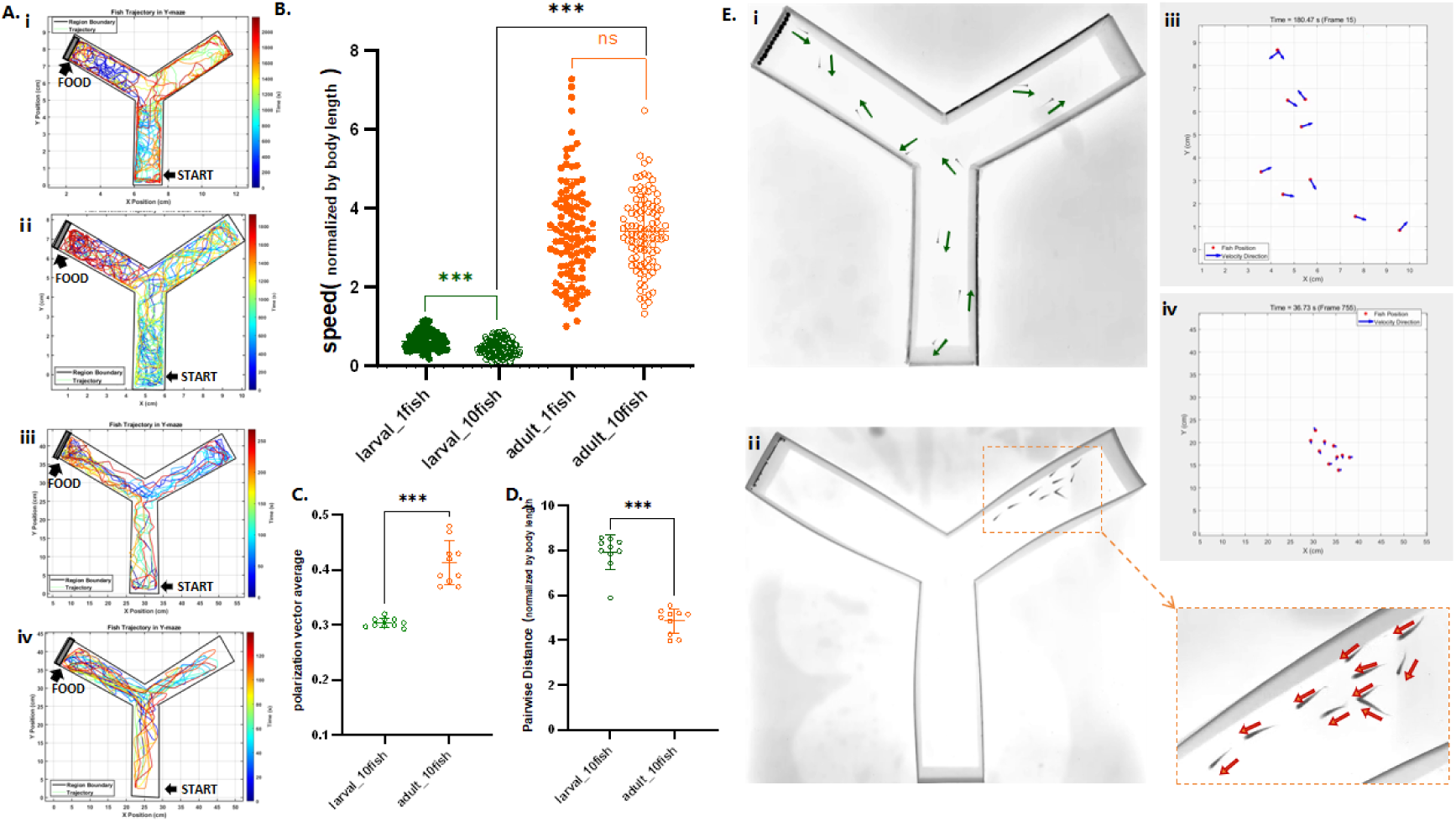
Dynamic analysis. **(A) Trajectories.** Representative movement trajectories of fish within the experimental arena, with colors representing different time points as indicated by the jet colormap on the right: colors ranging from blue (start of trial) to red (end of trial). (ⅰ) larvae, solitary; (ⅱ) larvae, group; (ⅲ) Adult, solitary; (ⅳ) Adult, group. **(B) Polarization vector average (PV).** T- test (two-tailed one-sample t-test) was used to examine differences between experimental groups. Hollow green dots represent the larvae group, and hollow orange dots represent the adult group. PV measures the degree of directional alignment among individuals, with higher values indicating greater polarization (i.e., more synchronized movement directions). p < 0.0001 indicates that polarization in adults was significantly higher than in larvae, ****. **(C) Pairwise distance (PD)** normalized by body length. PD measures the average distance between individuals after normalization by body length, with higher values indicating greater inter-individual distances and lower group cohesion. p < 0.0001 indicates that group cohesion in adults was significantly higher than in larvae, ****. **(D) Representative photographs** of (ⅰ) larvae group and (ⅱ) adult group, along with corresponding schematic diagrams illustrating movement directions for (ⅲ) larvae group and (ⅳ) adult group; arrows indicate movement direction. **(E) Swimming speed normalized by body length.** T-test (two-tailed one-sample t-test) was used to examine differences between experimental groups. Solid green dots represent the solitary larvae group, hollow green dots represent the larvae group, solid orange dots represent the solitary adult group, and hollow orange dots represent the adult group. larvae solitary vs larvae group: p < 0.0001, ****; adult solitary vs. adult group: p = 0.8399, p > 0.05, no correlation; larvae solitary vs. adult solitary: p < 0.0001, ****; larvae group vs. adult group: p < 0.0001, ****.

We next examined movement speed and found marked differences between adults and larvae, despite the similarity in spatial trajectories. After normalizing speed by body length, adults exhibited significantly higher overall swimming speeds than larvae (**Fig. 2B**; p < 0.0001), which partly explains the shorter foraging latencies observed at the individual level. Crucially, however, the effect of social condition on speed differed by age: in larvae, group individuals were significantly slower than solitary ones (p<0.0001), whereas in adults, no significant difference was observed between solitary and group conditions (p=0.8399). This indicates that speed alone cannot account for the group foraging advantage—adult groups did not forage more efficiently simply by swimming faster.

If speed is not the determining factor, how do adult groups achieve higher foraging efficiency? We therefore quantified two classic metrics of group structure: polarization vector average (PV), which measures directional alignment among individuals (higher values indicate more synchronized movement), and pairwise distance (PD), which measures group cohesion (lower values indicate tighter aggregation).

Our results revealed marked differences between adult and larvae groups on both metrics. Adult groups exhibited significantly higher PV values than larvae groups (**Fig. 2C**; p<0.0001), indicating more aligned movement directions. Concurrently, adult groups showed significantly lower PD values than larvae groups (**Fig. 2D**; p<0.0001), reflecting closer inter-individual distances and tighter aggregation. These quantitative findings reveal a critical distinction: adult groups display genuine social coordination, whereas larvae groups represent little more than a mere aggregation of individuals.

This conclusion was further supported by qualitative observations. As illustrated in **Fig. 2E** and **Movie 4**, larvae individuals were largely dispersed throughout the arena, with little apparent interaction among group members. In contrast, adult groups exhibited pronounced aggregation and coordinated movement, characterized by clear social interactions. Thus, the foraging advantage of adult groups stems from their heightened social coordination—a capacity that enables more efficient localization and approach to food resources. Larvae, which have not yet developed mature social coordination, fail to gain such foraging benefits even when tested in groups.

### Behavioral Interpretation – Age Differences in Stress Response and Social Buffering

The kinematic analyses presented above demonstrate that the foraging advantage of adult groups stems from enhanced social coordination. Why, then, do larvae groups fail to achieve similar coordination? To address this question, we examined behavioral factors, focusing on individual stress responses to a novel environment and their modulation by social context.

During speed analysis, we observed that some individuals exhibited prolonged periods of near-zero velocity—a characteristic freezing behavior indicative of stress. To minimize noise in the data, freezing was defined as events where swimming speed fell below 0.2 cm/s for larvae (or 1.0 cm/s for adults) and persisted for more than 2 s. **Fig.3A** shows representative speed traces (with freezing episodes highlighted in pink) for the four experimental conditions: solitary larvae (ⅰ), larvae group (ⅱ), solitary adult (ⅲ), and adult group (ⅳ). Visual inspection revealed that freezing occurred more frequently in larvae and was distributed throughout the trial, whereas freezing episodes in adults were relatively rare and brief.

**Figure 3.**
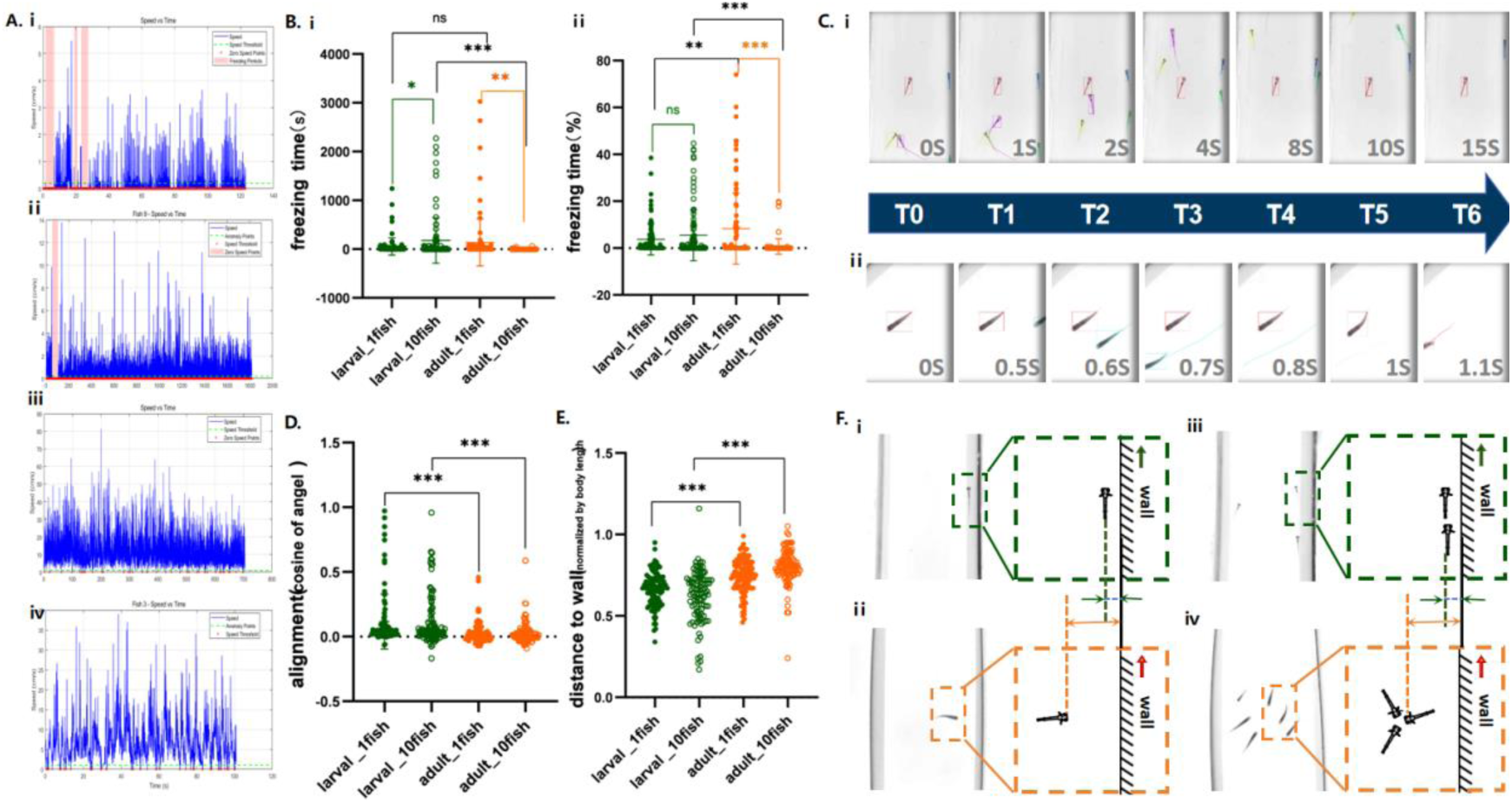
Behavioral analysis. **(A) Speed over time.** Blue lines represent the instantaneous swimming speed of fish throughout the trial. Red circles indicate time points where speed was zero; to account for noise in the data, zero speed was defined as < 0.2 cm/s for larvae and < 1.0 cm/s for adults. Pink shaded areas indicate freezing events. (ⅰ) larvae, solitary; (ⅱ) larvae, group; (ⅲ) Adult, solitary; (ⅳ) Adult, group. **(B)** (ⅰ) Freezing events were defined as episodes where swimming speed fell below 0.2 cm/s for larvae (or 1.0 cm/s for adults) and persisted for more than 2 s. Freezing time represents the cumulative duration of all freezing episodes throughout the trial. (ⅱ) Freezing % represents the proportion of freezing time relative to total trial duration. T-test (two-tailed one- sample t-test) was used to examine differences between experimental groups. Solid green dots represent the solitary larvae group, hollow green dots represent the larvae group, solid orange dots represent the solitary adult group, and hollow orange dots represent the adult group. (ⅰ) Freezing time results: larvae solitary vs. larvae group, p = 0.0149, p < 0.05, *; adult solitary vs. adult group, p = 0.0053, p < 0.05,**; larvae solitary vs. adult solitary, p = 0.1085, p > 0.05, no correlation; larvae group vs. adult group, p = 0.0002, p < 0.05, ***. (ⅱ) Freezing % results: larvae solitary vs. larvae group, p = 0.1794, p > 0.05, no correlation; adult solitary vs. adult group, p < 0.0001, ****; larvae solitary vs. adult solitary, p = 0.0065, p < 0.05, **; larvae group vs. adult group, p < 0.0001, ****. **(C) Time-series traces of behavioral responses.** Behavioral changes in (ⅰ) larvae and (ⅱ) adult fish when a conspecific passed nearby during a freezing episode. larvae remained frozen throughout the 15s period despite multiple conspecifics passing within their visual field. In contrast, adults responded almost instantaneously to passing conspecifics and exited the freezing state. **(D) Alignment (cosine of angle).** Alignment quantifies the degree to which a fish’s movement direction corresponds with the optimal direction (defined as the path along the wall toward the food patch). Values closer to 1 indicate higher alignment with the optimal direction; 0 indicates perpendicular movement; −1 indicates movement directly opposite to the optimal direction. Larvae solitary vs. adult solitary: p < 0.0001, ****; larvae group vs. adult group: p < 0.0001, ****. **(E) Distance to wall (normalized by body length).** Distance to wall measures the average distance between fish and the channel wall after normalization by body length, with higher values indicating greater distance from the wall. larvae solitary vs. adult solitary: p < 0.0001, ****; larvae group vs. adult group: p < 0.0001, correlation ****. **(F) Representative photographs and corresponding schematics** of (ⅰ) larvae solitary, (ⅱ) adult solitary, (ⅲ) larvae group, and (ⅳ) adult group. larvae consistently swam closer to the walls and spent most of their time moving along the walls. In contrast, adults were less influenced by walls during swimming; their movement was primarily driven by interactions with conspecifics, actively orienting toward group members and maintaining distance from walls.

Quantitative analysis confirmed this observation (**Fig.3B**). Under solitary conditions, no significant differences were detected between larvae and adults in total freezing duration (p = 0.1085), indicating that freezing is a common stress response to novel environments in zebrafish rather than an age-specific trait. However, the presence of conspecifics had markedly different effects on adults and larvae: adult groups showed significant reductions in both freezing duration(p = 0.0053) and percentage compared to solitary adults (p<0.0001), whereas larvae groups exhibited no such reduction—indeed, a trend toward increased freezing was observed. These results demonstrate that social interaction effectively buffers stress responses in adults but confers no such buffering effect in larvae. This provides an explanation for the absence of group foraging benefits in larvae: the presence of conspecifics fails to alleviate their stress, thereby preventing the establishment of coordinated group behavior.

Further analysis of individual behavioral responses during freezing episodes revealed the specific nature of this age difference. As illustrated in the representative time-series traces (**Fig. 3C**), when a freezing individual was approached or passed by a conspecific, larvae and adults responded in markedly different ways: larvae largely remained frozen, showing minimal interaction with the passing individual, whereas adults rapidly joined the moving conspecific and exited the freezing state (**Movie 5**). This observation indicates that adults can use social information to overcome stress-induced immobility—a capacity that larvae have not yet developed. This provides a direct behavioral mechanism for the adult group foraging advantage: conspecifics serve not only as partners in coordination but also as social resources for stress alleviation.

In addition to freezing behavior, larvae also exhibited distinctive patterns of spatial utilization. We analyzed the alignment between each fish’s movement direction and the “optimal foraging direction” (defined as the direction along the wall toward the food patch). Alignment was quantified as the cosine of the angle between the movement direction and the optimal direction, with values closer to 1 indicating higher alignment. Surprisingly, despite their lower foraging efficiency, larvae showed significantly higher alignment values than adults (**Fig. 3D**; p<0.0001), indicating that larvae were more likely to move along the optimal direction. This seemingly counterintuitive finding suggests that larvae wall-following behavior may reflect factors beyond physical models.

Analysis of the distance between fish and the channel wall (normalized by body length) revealed that larvae consistently stayed closer to the walls than adults (**Fig. 3E**; p<0.0001). Adults predominantly occupied the mid-channel region, whereas larvae were frequently observed near the walls (**Fig. 3F**). This spatial distribution difference is also evident in the trajectories shown in **Fig. 2A**: larvae paths trace clear outlines along the walls, whereas adult trajectories are more concentrated in the channel interior without pronounced boundary-following.

Based on these observations, we propose the following interpretation: owing to developmental immaturity (particularly incomplete swim bladder development), larvae may rely on physical contact with the wall for postural control and swimming stability, resulting in a strong edge preference. This edge preference accordingly aligns their movement direction with the wall-following path to the food, but such alignment reflects developmental constraints beyond physical models. In contrast, adults, with fully developed motor control, have no such reliance on walls; their movement direction is therefore more influenced by social factors (e.g., following conspecifics) than by mechanical orientation toward the food patch.

In summary, the failure of larvae groups to gain a foraging advantage can be attributed to two interrelated factors: (1) larvae exhibit heightened stress responses (frequent freezing) and lack social buffering from conspecifics; and (2) larvae edge preference stems from developmental constraints rather than beyond physical models, representing a form of pseudo-alignment that masks their true foraging incompetence. Together, these behavioral characteristics preclude the emergence of effective social coordination in larvae groups.

### Visual Social Cues Are Sufficient to Trigger Behavioral Benefits

The above results indicate that social interaction is the core driver of the group foraging advantage in adult zebrafish. However, does this interaction necessarily depend on real conspecifics, or is mere visual “social presence” sufficient to produce a similar effect? To address this question, we designed a mirror test: the inner walls of the channel were replaced with mirrors to provide the fish with a virtual visual companion (**Fig. 4A**). It should be noted that, due to material constraints, mirror-finish reflective paper (thickness < 0.1 mm, attached to PP plates) was used for larvae experiments, whereas acrylic mirrors were used for adult experiments; however, the reflective effect in water did not differ significantly between the two materials (**Fig. S3**). All other experimental parameters were consistent with those described previously.

**Figure 4.**
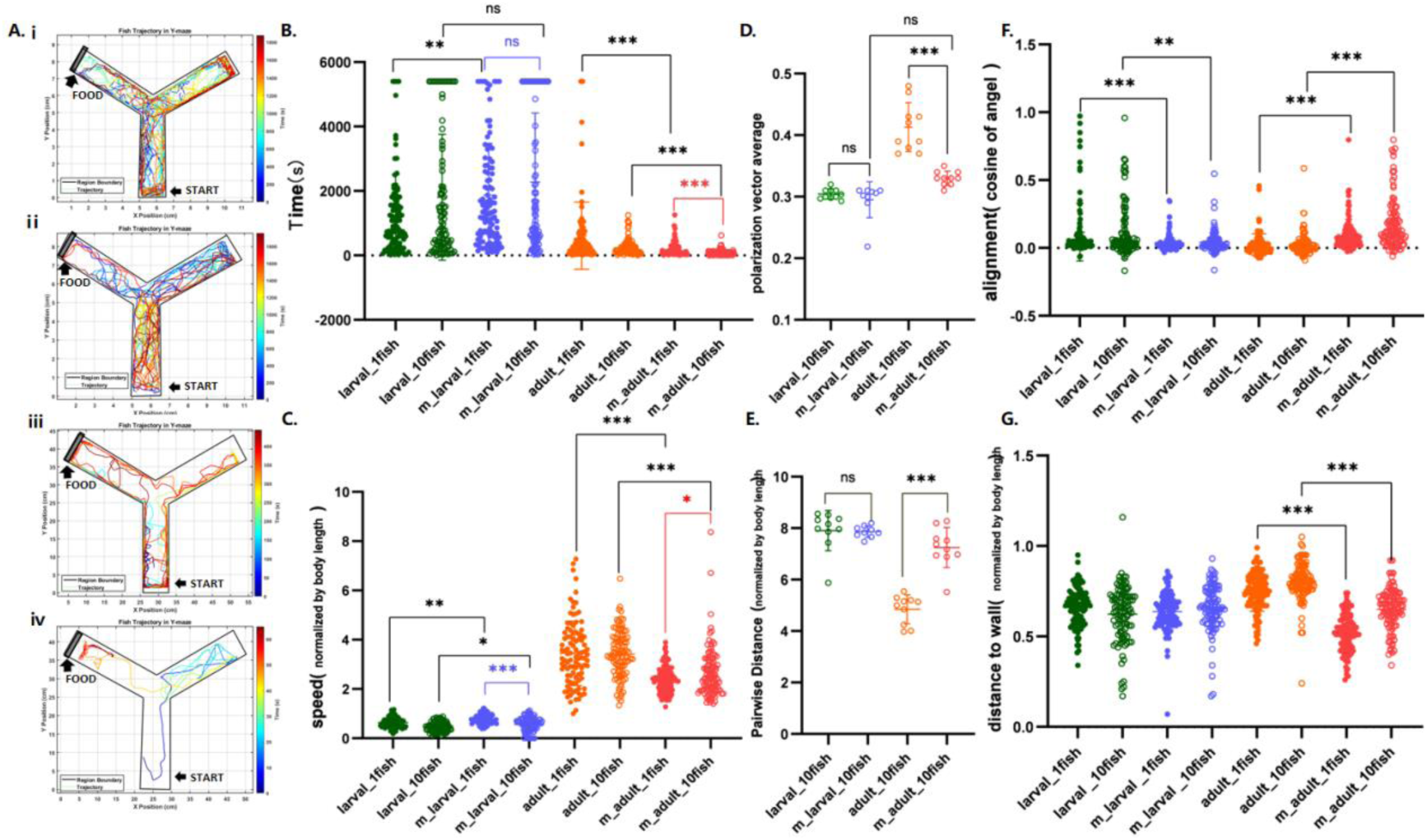
Mirror experiment. **(A) Trajectories.** Representative movement trajectories of fish within the experimental arena under mirror conditions, with colors representing different time points as indicated by the jet colormap on the right: colors ranging from blue (start of trial) to red (end of trial). (ⅰ) larvae, solitary; (ⅱ) larvae, group; (ⅲ) Adult, solitary; (ⅳ) Adult, group. **(B) Comparison of foraging time with and without mirror presentation.** T-test (two-tailed one-sample t-test) was used to examine differences between experimental groups. Solid green dots represent the solitary larvae group without mirror, hollow green dots represent the larvae group without mirror, solid orange dots represent the solitary adult group without mirror, and hollow orange dots represent the adult group without mirror; solid blue dots represent the solitary larvae group with mirror, hollow blue dots represent the larvae group with mirror, solid red dots represent the solitary adult group with mirror, and hollow red dots represent the adult group with mirror. Solitary larvae without mirror vs. solitary larvae with mirror: p = 0.0025, p < 0.05, correlation **; larvae group without mirror vs. larvae group with mirror: p = 0.1036, p > 0.05, no correlation; solitary adult without mirror vs. solitary adult with mirror: p = 0.0002, p < 0.05, correlation ***; adult group without mirror vs. adult group with mirror: p < 0.0001, correlation ****. **(C) Swimming speed normalized by body length.** Solitary larvae with mirror vs. larvae group with mirror: p < 0.0001, correlation ****; solitary adult with mirror vs. adult group with mirror: p = 0.0116, p < 0.05, correlation *; solitary adult without mirror vs. solitary adult with mirror: p < 0.0001, correlation ****; adult group without mirror vs. adult group with mirror: p < 0.0001, correlation ****. **(D) Polarization vector average (PV)**. PV measures the degree of directional alignment among individuals, with higher values indicating greater polarization (i.e., more synchronized movement directions). Adult group without mirror vs. adult group with mirror: p < 0.0001, correlation ****. **(E) Pairwise distance (PD) normalized by body length.** PD measures the average distance between individuals after normalization by body length, with higher values indicating greater inter-individual distances and lower group cohesion. Adult group without mirror vs. adult group with mirror: p < 0.0001, correlation ****. **(F) Alignment (cosine of angle).** Alignment quantifies the degree to which a fish’s movement direction corresponds with the optimal direction (defined as the path along the wall toward the food patch). Values closer to 1 indicate higher alignment with the optimal direction; 0 indicates perpendicular movement; −1 indicates movement directly opposite to the optimal direction. Adult group without mirror vs. adult group with mirror: p < 0.0001, correlation ****. **(G) Distance to wall (normalized by body length).** Distance to wall measures the average distance between fish and the channel wall after normalization by body length, with higher values indicating greater distance from the wall. Adult group without mirror vs. adult group with mirror: p < 0.0001, correlation ****.

The presence of the mirror induced marked changes in fish movement trajectories. As shown in **Fig. 4A**, fish in all four conditions—solitary larvae (ⅰ), larvae group (ⅱ), solitary adult (ⅲ), and adult group (ⅳ)—exhibited a tendency to swim closer to the mirror side, with trajectories forming clear linear boundaries along the mirror. This phenomenon was particularly pronounced in adults: adults swam distinctly along the mirror surface, displaying typical “parallel swimming” behavior. This suggests that fish may perceive their mirror image as a conspecific, thereby eliciting social approach behavior similar to that observed in real groups.

Furthermore, the mirror condition significantly enhanced foraging efficiency. As shown in **Fig. 4B**, the presence of the mirror produced divergent effects across ages and social conditions. For larvae, it significantly increased the foraging latency of solitary individuals (p = 0.0025), but exerted no significant influence on the larval group (p = 0.1036). In contrast, for adults, the mirror significantly reduced the latency in both solitary (p = 0.0002) and group conditions (p < 0.0001). Notably, even with the mirror present, the adult group still displayed a shorter foraging latency than solitary adults (p < 0.0001). This finding is highly consistent with the results obtained from real groups, providing further evidence that the facilitative effect of social cues—even virtual ones—on foraging efficiency depends on the individual’s developmental stage.

We further analyzed kinematic parameters under the mirror condition. Unexpectedly, despite the mirror-induced improvement in foraging efficiency, swimming speed did not increase; rather, it decreased significantly. As shown in **Fig. 4C**, adults in particular exhibited significantly lower swimming speeds under the mirror condition compared to the no-mirror control (p < 0.0001), and the inter-individual variability in speed within groups was also reduced. Concurrently, the presence of the mirror significantly reduced the frequency and duration of freezing behavior (**Fig. S5B**).

Integrating these results, we propose the following interpretation: the visual social cues provided by the mirror create a perception of “being accompanied,” which reduces the fish’s anxiety levels, resulting in smoother locomotion with fewer stress-induced interruptions. This calming effect, rather than an increase in speed, may represent the underlying mechanism by which the mirror promotes foraging efficiency.

A more intriguing finding emerged from the “anomalous” changes in group parameters. In the real-group experiments, adult groups exhibited high polarization (PV) and low inter-individual distance (PD)—hallmarks of effective social interaction. However, under the mirror condition, adult groups showed significantly reduced PV (**Fig. 4D**; p < 0.0001) and increased PD (**Fig. 4E**; p < 0.0001), with values regressing to levels comparable to those of larvae groups. At first glance, this result appears paradoxical: if the mirror enhances foraging efficiency, why does it simultaneously disrupt group coordination?

Our interpretation is as follows: the mirror alters the target of social interaction. Under mirror-free conditions, adults interact with real conspecifics, forming highly coordinated groups. Under mirror conditions, however, fish shift their attention to their mirror image—each fish interacts with its own reflection rather than with real conspecifics (**Movie 6**). From the experimenter’s perspective, the coordination among real fish (as measured by PV and PD) naturally declines because each individual’s “social attention” is diverted toward its mirror image. Nevertheless, this interaction with the mirror image itself constitutes a form of “quasi-social interaction” sufficient to produce calming and behavioral facilitating effects.

This interpretation is supported by other parameters. Under the mirror condition, adult alignment (with respect to the optimal foraging direction) increased significantly (p < 0.0001), while distance to the wall decreased significantly (p < 0.0001)—that is, adults exhibited edge preference similar to that of larvae. This indicates that under mirror conditions, adults swim more along the wall—not due to developmental constraints (as in larvae, which may require wall contact for postural support), but because the mirrored wall becomes the focal point of social interaction.

In summary, the mirror test provides three lines of evidence supporting the critical role of social interaction in foraging efficiency. First, visual social cues alone are sufficient to produce behavioral benefits: even in the absence of real conspecifics, a mere mirror image significantly enhances foraging efficiency, reduces freezing behavior, and stabilizes swimming speed. This suggests that the “feeling of being accompanied” has a calming effect sufficient to improve foraging performance. Second, this effect is age-dependent: adults derive greater benefits from the mirror than larvae, consistent with the results from real-group experiments and further confirming developmental differences in social responsiveness. Third, the mirror test reveals multiple dimensions of social interaction: efficient foraging in real groups depends on group coordination (high PV, low PD), whereas efficient foraging under mirror conditions depends on emotional calming (speed stabilization, reduced freezing). Both mechanisms may operate in real groups—the presence of conspecifics provides both the “instrumental value” of coordinated cooperation and the “social support value” of emotional buffering. Adults are capable of fully utilizing both benefits, whereas larvae possess neither.

## Summary and Discussion

This study investigated how group behavior influences foraging efficiency in zebrafish and how this influence depends on developmental stage. By comparing the foraging performance of larvae and adults in both solitary and group contexts, we found that grouping significantly enhanced foraging efficiency in adults but provided no such advantage for larvae. This difference stems from the nature of the groups themselves: adults formed highly coordinated social groups characterized by higher polarization (PV) and smaller inter-individual distances (PD), whereas larval groups were merely simple aggregations lacking effective social interaction. Mechanistically, the group advantage in adults operates on two levels. At the kinematic level, social coordination enabled efficient spatial exploration and information transfer. At the behavioral level, adults benefited from “social buffering” by conspecifics, which significantly reduced stress responses (e.g., freezing behavior) in a novel environment. Larvae, in contrast, not only exhibited higher baseline stress but also failed to receive this emotional buffering from their peers. Mirror experiments further revealed that even purely visual social cues (mirror images) were sufficient to enhance foraging efficiency via emotional buffering. However, this effect was also age-dependent and could not substitute for the coordinated interactions observed in genuine groups.

In summary, this study reveals a dual mechanism through which group behavior promotes foraging efficiency—the instrumental value of social coordination and the emotional value of social buffering—and demonstrates that the realization of both benefits is contingent upon developmental maturation. In a more general perspective, individuals need to become mature enough to spontaneously execute sophisticated social and group behaviors at a higher level [44]. These findings offer new insights into the developmental trajectory of social behavior. Future research integrating long-term observations, brain and neural activity [46–47], and gene editing technologies will be essential to elucidate the neural foundations and critical windows for the development of social competence.

## Materials and Methods

### Experimental Subjects and Rearing Conditions

The zebrafish used in this experiment were first-generation offspring obtained through natural spawning from wild-type broodstock purchased from a professional supplier, after a two-week acclimatization period in the laboratory. The housing and maintenance of experimental zebrafish followed the methods described by Avdesh et al [36,37]. Post-hatching larvae were temporarily housed in static aquaria outside the recirculating system, with daily water changes. The water temperature was constantly maintained at 26.0 ± 0.5°C, pH at 7.0–7.5, and conductivity at 500 ± 50 µS/cm. A fixed photoperiod of 14 hours light / 10 hours dark was implemented. Larvae were fed commercial starter diets (Zeigler Ⅰ and Ⅱ) daily until 16 days post-fertilization (dpf). After 16 dpf, they were transferred to the recirculating system and fed daily with commercial feed and freshly hatched Artemia nauplii.

Larvae used in this experiment were 10 dpf old, with a body length of approximately 0.5 cm; adult fish were 80 dpf old, with a body length of approximately 2.5 cm [37]. All experimental individuals were healthy, age-matched, and size-matched. They were fasted for 24 hours prior to the experiment to eliminate the influence of feeding status on behavioral testing.

### Experimental Apparatus

A custom-built video tracking system was employed for this experiment. Following common practices for recording zebrafish behavioral data, an industrial camera was mounted perpendicularly above the experimental tank [32]. A surface light source placed at the bottom of the tank served as the sole illumination to provide a consistent and uniform lighting environment (**Fig. 1A**). The experimental tanks were rectangular parallelepipeds made from 1 mm thick, semi-translucent white frosted PP board. Internal partitions made of the same material were added to create a Y-maze configuration. Two different tank sizes were established based on the developmental stage and body length of the experimental subjects, adapting methods from Giana de P. Cognato et al [45]. For larvae (10 dpf, body length ∼0.5 cm), the Y-maze arm length was 5.0 cm, corridor width 1.6 cm, height 1.6 cm, and water depth 1.5 cm. For adult fish (80 dpf, body length ∼2.5 cm), the Y-maze arm length was 25.0 cm, corridor width 8.0 cm, height 8.0 cm, and water depth 7.5 cm[32]. For the mirror test, the walls of the larval tank were lined with mirror-reflective paper (thickness < 0.1 mm), while the adult tank walls were constructed using acrylic mirrors to create a mirror reflection effect.

### Experimental Design

Experiments were conducted during the zebrafish active period (10:00–19:00 daily) [32]. Two hours prior to the test, the experimental fish, along with their home tanks, were transferred to the testing room to minimize the impact of handling stress. An oxygen pump was provided to prevent hypoxia. The testing environment was an open, temperature-controlled room ensuring ample oxygen and stable temperature. During the test, the surface light source was placed horizontally on an anti-vibration table. The experimental tank was then placed on the light source, filled with fresh experimental water (26.0 ± 0.5°C).

Floating feed was first placed in the food arm of the Y-maze (in this experiment, the food location was fixed in the upper-left corner of the test area). Subsequently, a fish was gently introduced into the start arm. The entire foraging process was recorded using the overhead industrial camera (30 frames per second). Fish used in an experiment were not reused in subsequent trials to ensure all experimental subjects were naive to the apparatus, avoiding the influence of memory and learning [32]. The experiment comprised four groups: larval individual (n=1), larval group (n=10), adult individual (n=1), and adult group (n=10) [42,43].

### Behavioral Metrics and Definitions

Foraging efficiency was defined as the time interval from when the fish entered the test area until it first touched the food [39]. Polarization was calculated as the mean vector length of the group, with values closer to 1 indicating higher directional alignment [40,41]. Alignment was defined as the degree of alignment between an individual fish’s movement direction and the optimal direction. A value of 1 indicates perfect alignment, 0 indicates perpendicular movement (neutral), and −1 indicates movement in the completely opposite direction. The spatial distance between individuals within a group was quantified using the mean nearest neighbor distance, i.e., the mean of the Euclidean distances between all pairs of fish, following the method of Yang et al [40]. Freezing behavior was defined as events where larval swimming speed fell below 0.2 cm/s for more than 2 seconds, or adult speed fell below 1.0 cm/s for more than 2 seconds [17,18]. The duration and percentage of total time spent freezing were quantified. Edge preference was defined as the average distance of the fish from the tank wall (normalized by body length) [18].

### Video Tracking and Data Analysis

Trajectory tracking was performed using idtracker.ai, an algorithm based on deep learning and contrastive learning that enables high-precision multi-target tracking of unmarked animals, yielding the movement coordinates of each fish over time during foraging[38]. The exact frame of the first successful foraging event was identified through manual calibration. Subsequent data analysis was conducted using MATLAB (R2023a) to calculate behavioral parameters for each experimental group, including foraging latency, swimming speed, alignment, and freezing duration. Statistical analysis was performed using GraphPad Prism (8.0.2), with appropriate statistical tests applied for group comparisons. The extraction methods for behavioral parameters followed standard practices in zebrafish group behavior research.

## Acknowledgments

We thank Deepseek v2.2.1 for text polishing. This study was supported by taxpayers of China through National Natural Science Foundation of China (Distinguished Young Scholars Funding, Overseas; Young Scientists Fund - Category C) and Wuhan University (Talents Startup Funding).

## Author contributions

B.L. conceptualized the project. B.L. and P.S.W. designed the research. P.S.W. performed the experiment and analyzed the data with the help of M.Y.C.. All authors discussed the results and wrote the paper.

## Competing interests

The authors declare no competing interests.

### Additional information

**Supplementary figures** are available for this paper after the main text.

**Correspondence and requests for materials** should be addressed to B.L.

### Data availability

The data and analysis code that support the findings of this study available upon request to the correspondence author.

## Figure Captions

**Figure S1.**
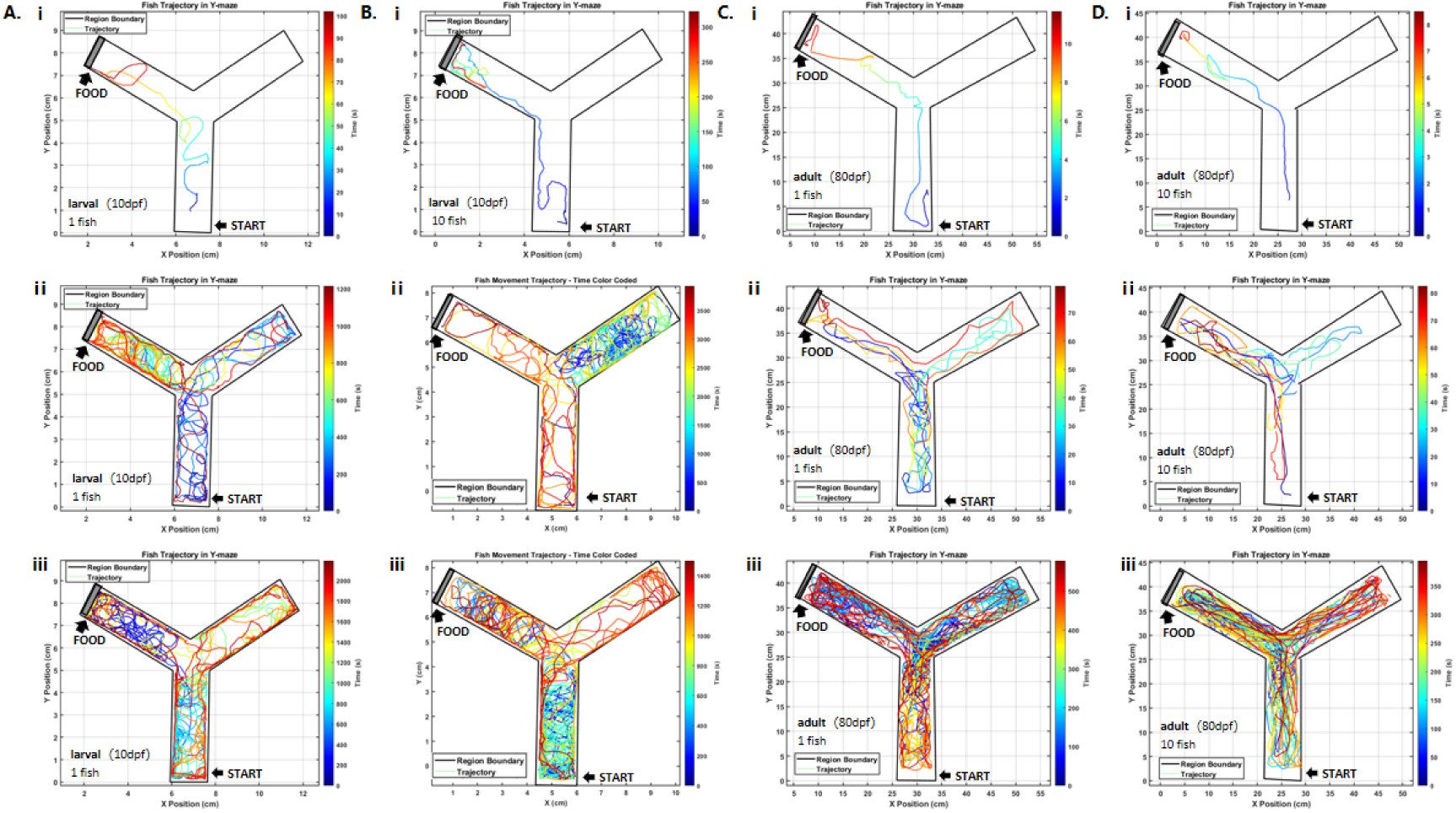
Trajectories. Representative movement trajectories of fish within the experimental arena without mirror conditions, with colors representing different time points as indicated by the jet colormap on the right: colors ranging from blue (start of trial) to red (end of trial). **(A)** larvae, individual; **(B)** larvae, group; **(C)** adult, individual; **(D)** adult, group.The starting point and the food location are marked in the figure. The black square frame indicates the boundary of the experimental arena, and the coloured lines represent the trajectories of the fish swimming within the arena. Different colours indicate different time points, as shown in the jet colour bar on the right: colours closer to blue indicate times nearer to the start of the experiment, while colours closer to red indicate times nearer to the end of the experiment. Exploratory behaviour shows certain individual differences. Although the vast majority of individuals choose to explore multiple times before deciding to forage, a small proportion of individuals swim directly towards and consume the food once they have detected it. This variability exists regardless of the individual’s developmental stage, group level, or social environment.

**Figure S2.**
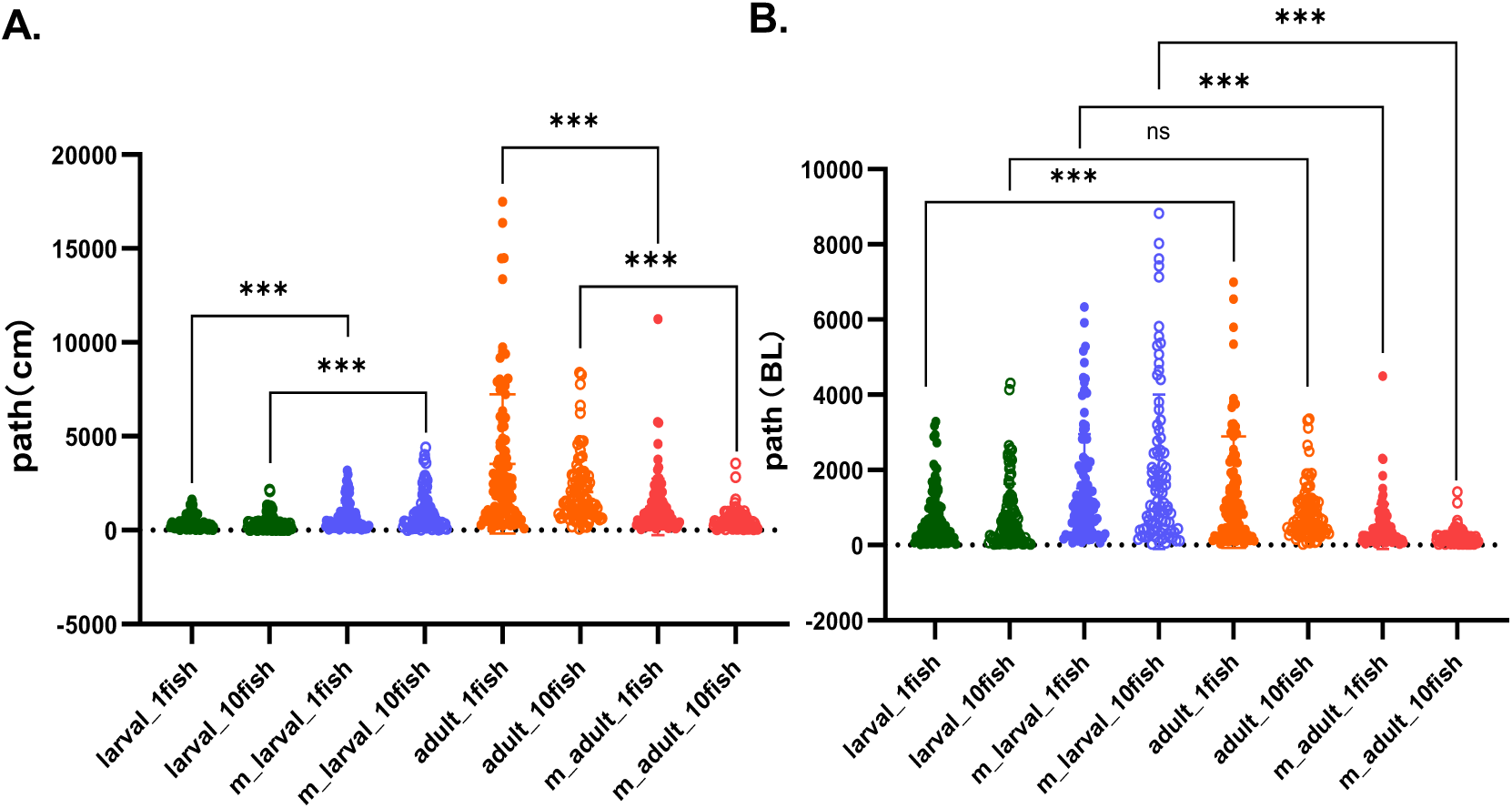
Comparison of total movement path lengths. **A. Path; B**. **Path normalized by body length**; larval BL=0.5cm; adult BL=2.5cm; T-test (two-tailed one-sample t-test) was used to examine differences between experimental groups. Solid green dots represent the individual larvae group without mirror, hollow green dots represent the larvae group without mirror, solid orange dots represent the individual adult group without mirror, and hollow orange dots represent the adult group without mirror; solid blue dots represent the individual larvae group with mirror, hollow blue dots represent the larvae group with mirror, solid red dots represent the individual adult group with mirror, and hollow red dots represent the adult group with mirror. In the absence of a mirror, after normalizing the path lengths by body length, there was no significant difference in the total movement path length between adult groups and larval groups (p > 0.005). The total path length of individual adults was even slightly longer than that of individual larval. The addition of a mirror had a significant effect on reducing the movement path length in adult fish, both in individual individuals (p < 0.0001, correlation ****) and in groups (p < 0.0001, correlation ****), with the total path length being markedly shortened after the mirror was introduced.

**Figure S3.**
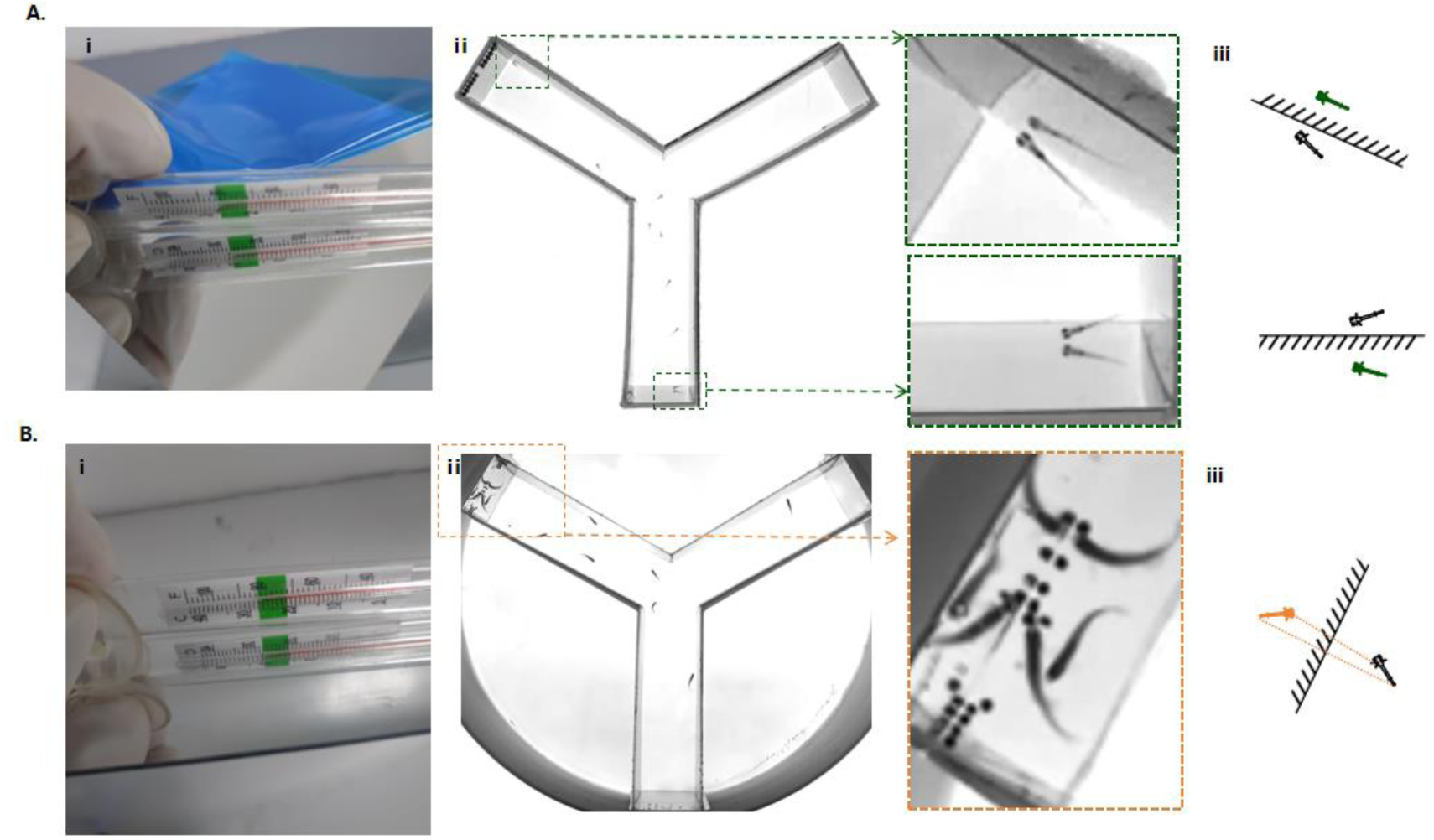
Materials used in the mirror experiment. **A.** For larval fish, the mirror material used was adhesive-backed reflective paper, with a thickness of less than 0.1 mm, attached tightly to the 1-mm-thick white polypropylene tank wall. **B.** For adult fish, the mirror material used was an acrylic mirror, with a thickness of 1 mm. The two materials produced similar effects in water, and all other experimental parameters were the same as previously described.

**Figure S4.**
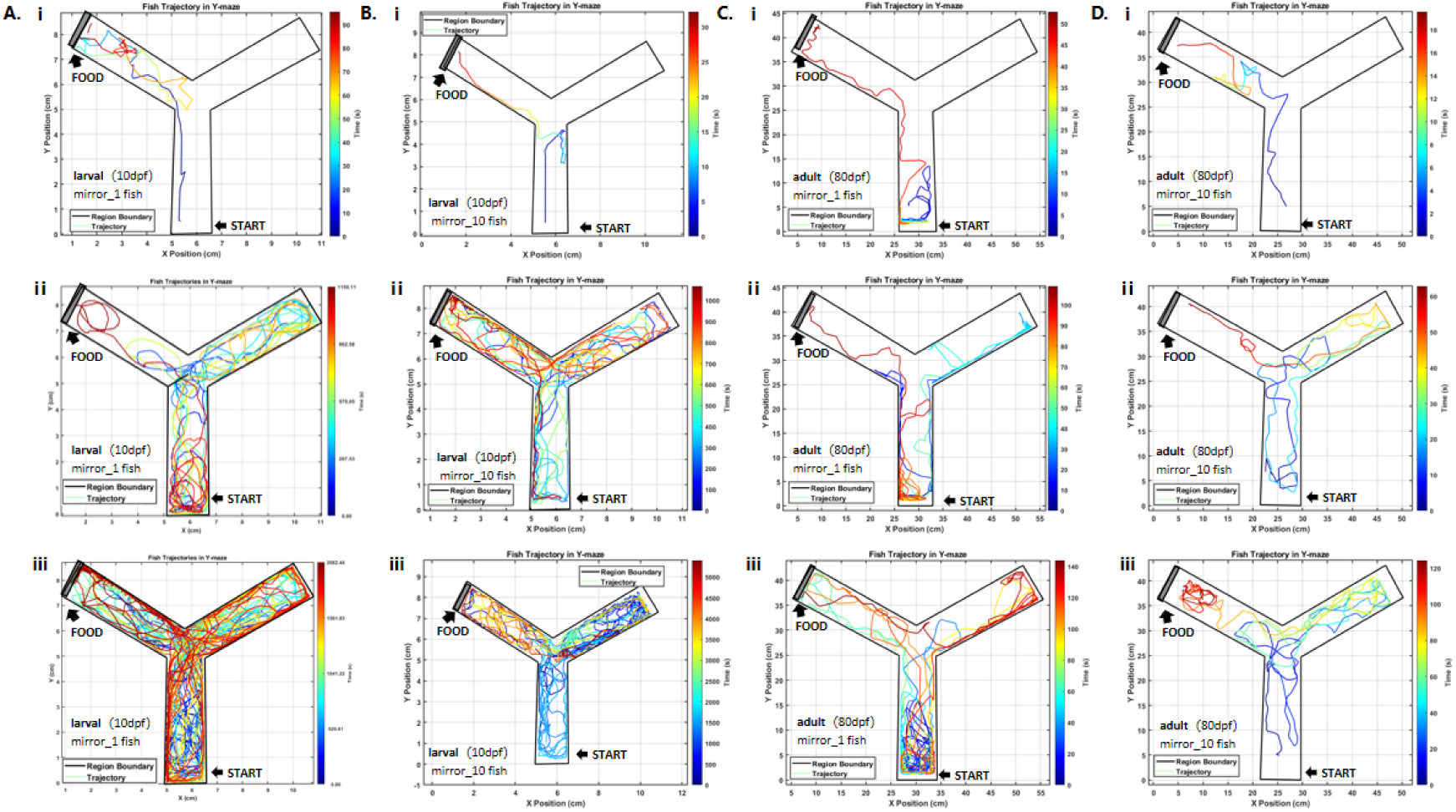
Trajectories. Representative movement trajectories of fish within the experimental arena under mirror conditions. (A) larvae, individual; (B) larvae, group; (C) adult, individual; (D) adult, group. The starting point and the food location are marked in the figure. The black square frame indicates the boundary of the experimental arena, and the coloured lines represent the trajectories of the fish swimming within the arena. Different colours indicate different time points, as shown in the jet colour bar on the right: colours closer to blue indicate times nearer to the start of the experiment, while colours closer to red indicate times nearer to the end of the experiment.

**Figure S5.**
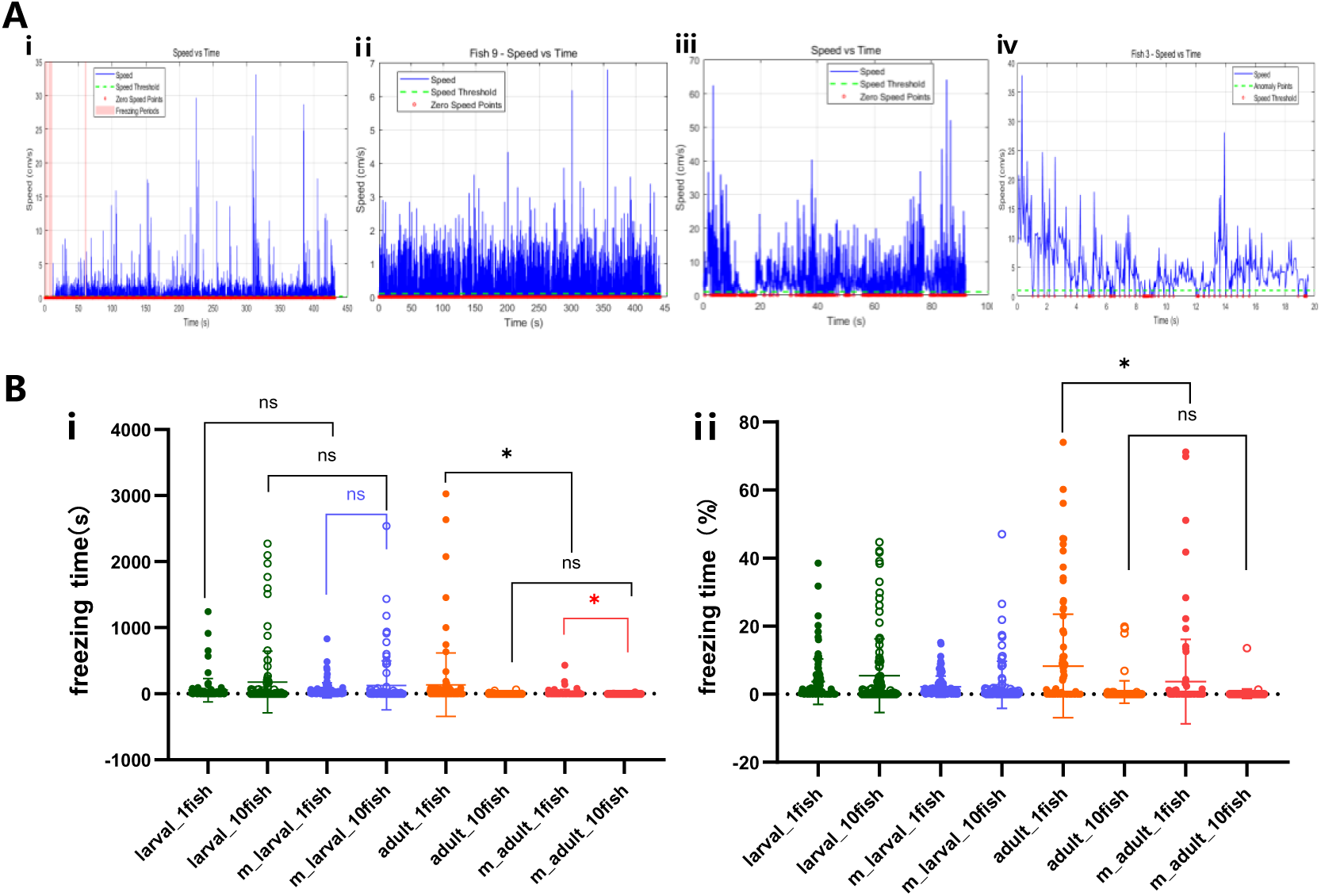
Behavioral analysis with mirror. **(A)** speed in time. The blue lines represent the curves of swimming speed over time for fish moving in a Y-maze composed of mirrors. Red circles mark the positions where the speed is zero. Considering noise in data processing, speeds below 0.02 cm/s for larval fish and below 0.1 cm/s for adults were defined as zero. Pink background areas indicate freezing events.(i) larval, individual; (ii) larval, group; (iii) adult, individual; (iv) adult, group.(B) (i) Freezing events were defined as swimming speeds below 0.2 cm/s for larval fish and below 1 cm/s for adults, both lasting for more than 2 consecutive seconds. Freezing time is the sum of all freezing episodes throughout the entire experiment. (ii) Freezing% represents the proportion of freezing time relative to the total movement time. The correlations between experimental groups were verified by t-test (two-tailed one-sample t-test). Solid green dots represent the larval individual group without a mirror, open green dots represent the larval group without a mirror, solid orange dots represent the adult individual group without a mirror, and open orange dots represent the adult group without a mirror. Solid blue dots represent the larval individual group with a mirror, open blue dots represent the larval group with a mirror, solid red dots represent the adult individual group with a mirror, and open red dots represent the adult group with a mirror. After the addition of the mirror, the freezing time of adult groups decreased further compared to that of individual adults (p = 0.0138, p < 0.05, *).

